# Gut microbial metabolites butyrate and acetate limit Zika virus replication and associated ocular manifestations via the G-protein coupled receptor 43/FFAR2

**DOI:** 10.1101/2025.07.15.664962

**Authors:** Nikhil Deshmukh, Prince Kumar, Lal Krishan Kumar, Vaishnavi Balendiran, Pawan Kumar Singh

## Abstract

Short-chain fatty acids (SCFAs) are gut microbial metabolites produced by gut microbiota from dietary fiber. SCFAs have shown both pro- and anti-viral roles among different viruses, and are known to regulate immune functions during infections. However, their role against the Zika virus (ZIKV) in general and ocular infection, in particular, has never been investigated. In the present study, we aimed to examine the role of three SCFA derivatives: phenylbutyrate (PBA), sodium butyrate (NaB), and sodium acetate (NaAC), on ZIKV replication and associated ocular complications using primary human trabecular meshwork cells (HTMCs) and an IFNAR1-deficient mouse model of ocular infection. Our findings reveal that PBA and NaAc treatment dramatically suppressed the ZIKV replication in HTMCs. NaB showed a slightly less effect than PBA and NaAc. PBA and NaAc treatment significantly attenuated the ZIKV-induced inflammatory cytokine, interferons, and interferon-stimulated genes response via antagonizing the RIG-I/NFκB/MAPKs/STAT1-3 signaling pathways. We discovered that ZIKV induces the expression of free fatty acid receptor 2 (FFAR2)/ GPR43 in HTMCs, which is further potentiated by PBA/NaAc. Pharmacological inhibition of FFAR2 abrogated the protective abilities of PBA/NaAc and significantly increased viral replication. Blocking FFAR2 receptors promoted ZIKV-induced cell death, which was suppressed by PBA and NaAc. Mechanistically, butyrate and acetate inhibited ZIKV binding and cellular entry and inactivated the virus before internalization. PBA and NaAc treatment in mice attenuated the ZIKV-induced ocular manifestations (intraocular pressure, RPE/retinal atrophy, and anterior segment inflammation), which was abrogated by FFAR2 inhibition. Collectively, our findings indicate that SCFA treatment is an effective approach to limit ZIKV replication and associated ocular damage and may be worth exploring as a means to treat or prevent ZIKV-induced ocular complications in humans.

**Importance:** ZIKV is known to cause severe ocular manifestations in *in-utero* exposed infants; however, the molecular mechanisms of ZIKV-induced ocular complications remain unknown. SCFAs have demonstrated both pro- and anti-viral roles against different viruses; however, their role against ZIKV is unknown. We showed that SCFAs butyrate and acetate suppress ZIKV transmission and associated ocular complications. The anti-ZIKV activity of these SFACs is mediated via FFAR2, and pharmacological inhibition of FFAR2 promotes ZIKV-induced inflammatory and cell death responses, as well as ocular malformations.

## Introduction

Zika virus (ZIKV) is a global public health threat due to its association with severe neurological and ocular manifestations collectively referred to as congenital Zika syndrome(1). ZIKV gained global attention in 2015 via an epidemic in the Americas that affected millions of people, resulting in the declaration of a Public Health Emergency of International Concern (PHEIC) by WHO(2–4). Although PHEIC was lifted in November 2016, ZIKV remains a concern due to its ongoing transmission in some geographical areas and potential long-term effects on affected individuals. Currently, ZIKV is circulating in >89 countries, and sporadic outbreaks are reported in different regions around the world. Therefore, constant surveillance is essential to ensure preparedness for early detection and treatment of future ZIKV pandemics(5, 6). ZIKV is a positive-sense RNA virus that belongs to the *Flaviviridae* family. It is an arbovirus and is primarily transmitted through mosquito bites. However, it can also spread via sexual intercourse, blood transfusion, organ transplantation, and maternal-fetal transmission(7, 8). *In utero* exposure to ZIKV is linked to severe microcephaly, vision and hearing impairment, as well as articular and musculoskeletal abnormalities(1, 9). Among ocular manifestations, ZIKV caused a wide range of ocular symptoms, including chorioretinal atrophy, hypoplasia, focal pigmented mottling, RPE mottling, retinal focal spots, severe retinal vessel attenuation, optic nerve atrophy, optic disc anomalies, and congenital glaucoma(10–16). However, the ocular disease pathogenesis and long-term consequences of ZIKV infection remain unknown. Besides, no specific antiviral therapies or vaccines are currently available for ZIKV; therefore, it is essential to investigate alternative antivirals to treat/prevent ZIKV-induced complications.

Short-chain fatty acids (SCFAs) such as butyrate, propionate, and acetate are gut microbial metabolites that are formed from colonic fermentation of dietary fiber. These metabolites maintain gut barrier integrity and are also known to modulate host immunity and pathogen susceptibility(17–20). SCFAs act by inhibiting either class I and IIa histone deacetylases (HDACs) or by binding to free fatty acid receptors (FFAR2 or FFAR3)(17) (18). FFARs belong to the G-protein coupled receptors (GPCR) family, such as GPR43 (FFAR2) and GPR41 (FFAR3). SCFAs act as ligands for FFARs and, upon activation, regulate multiple signaling pathways including ERK1/2, p38, JNK, Akt, intracellular Ca^+2^ release, and inhibition of cAMP accumulation. Recent studies have shown that both SCFAs and their receptors FFARs, play a critical role in the regulation of inflammation and adaptive immunity(18–22). They are also known to modulate the pathogenesis of various neurological and ocular diseases(18, 23–27). SCFAs have shown differential roles against invading microbial pathogens, including bacteria and viruses(27–29). Among viruses, SCFA butyrate has been shown to promote the replication of influenza A virus (IAV), human immunodeficiency virus 1 (HIV-1), human metapneumovirus (hMPV), and vesicular stomatitis virus (VSV)(30, 31). In contrast, SCFA treatment has been suggested to suppress viral replication and associated inflammation in SARS-CoV-2(22), Herpes simplex virus 1 (HSV-1) (32–34), and hepatitis B virus (HBV)(35) infections. However, the role of SCFAs and their receptor FFAR2 in ZIKV infection and associated inflammatory immune responses has never been demonstrated.

In the present study, we reported the previously unknown role of SCFA derivatives phenylbutyrate (PBA), sodium butyrate (NaB), and sodium acetate (NaAc), and a GPCR-FFAR2 in ZIKV ocular infection. Our data show that butyrate (PBA, NaB) and acetate (NaAc) derivatives suppressed ZIKV replication and associated ocular manifestations. PBA and NaAc diminished ZIKV-induced inflammatory, interferons (IFNs), and interferon-stimulated gene (ISGs) response. Their activity was mediated via FFAR2, and pharmacological inhibition of FFAR2 significantly enhanced the viral replication, associated cell death, and ocular complications.

## Results

### SCFAs phenylbutyrate and sodium acetate restrict ZIKV replication in HTMC

SCFAs butyrate and acetate have demonstrated differential antiviral activities against different viruses. It has been reported to inhibit HSV-1 replication and associated ocular inflammation(32–34) while promoting IA, HIV-1, hMPV, and VSV, and has no effect on the Sendai virus (SeV) replication(30, 31). However, their role in flaviviruses, such as ZIKV infection, is entirely unknown. Here, we investigated the role of three SCFAs: PBA, NaB, and NaAc in ZIKV replication using human primary trabecular meshwork cells (HTMCs). Previously, we demonstrated that ZIKV has high tropism towards the anterior segment of the eye and can permissively infect HTMCs(11). To investigate the role of these SCFAs, we first pretreated HTMC with PBA, NaB, and NaAc prior to ZIKV infection. We observed that PBA and NaAc significantly inhibited the ZIKV replication as measured by ZIKV-envelope (E) antigen (4G2) immunofluorescence staining (**Fig. 1A**). To further confirm the inhibition of ZIKV replication by these SCFAs, we performed immunoblotting for ZIKV nonstructural protein NS3. Our data show that PBA and NaAc significantly inhibited ZIKV replication, as indicated by the suppression of NS3 protein in treated cells compared to untreated cells (**Fig. 1B&C**). NaB also demonstrated ZIKV inhibitory properties, although to a slightly lesser extent than PBA and NaAc (**Fig. 1A-C**). To further examine the effect of butyrate and acetate on the number of replicating virions, we performed a plaque assay. Our results revealed significant inhibition of ZIKV plaques by PBA, NaB, and NaAc compared to untreated cells, confirming their anti-ZIKV role (**Fig. 1D**). Together, our findings confirm that SCFAs phenylbutyrate and sodium acetate can inhibit ZIKV replication and transmission.

**Figure 1:**
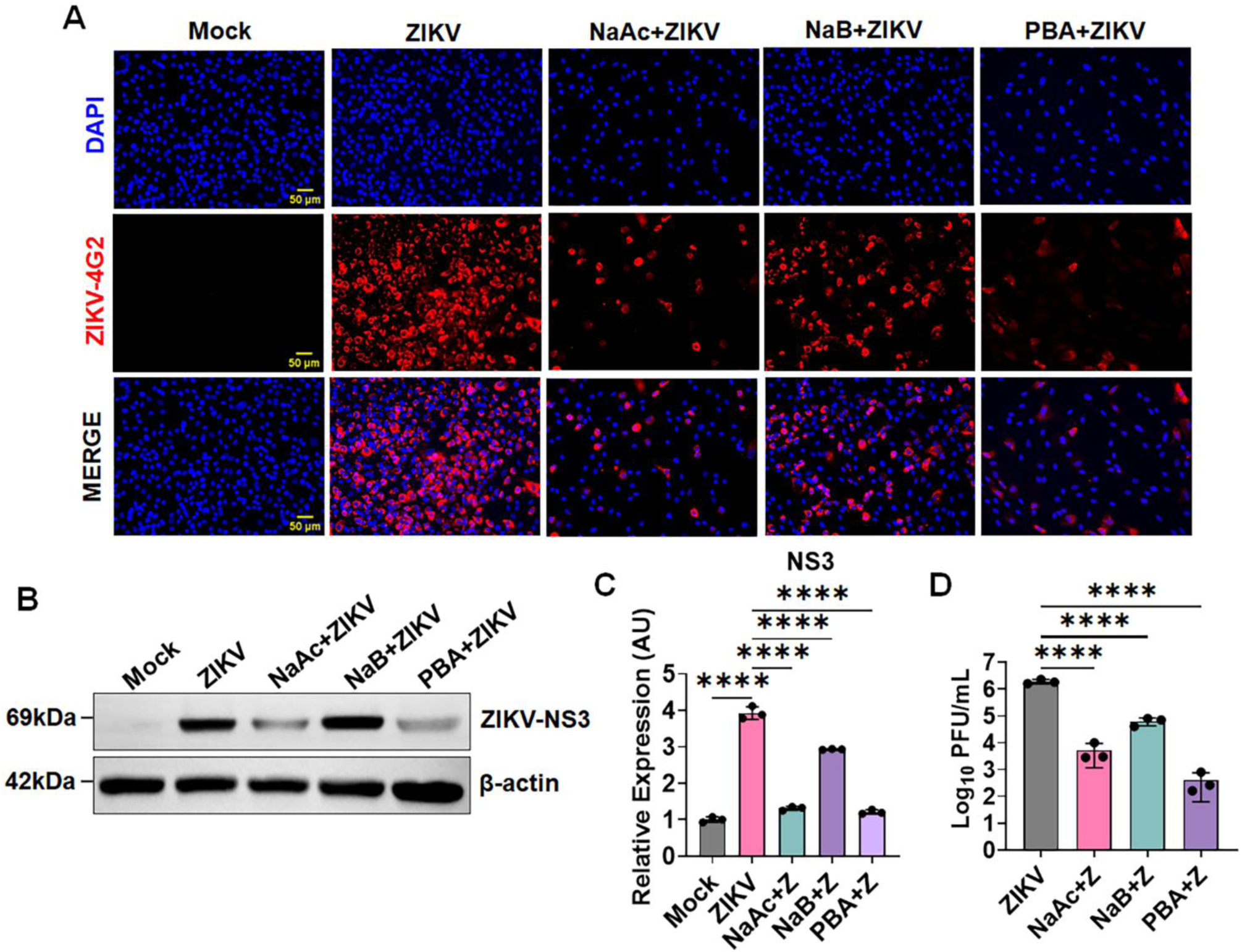
SCFA restricts ZIKV replication in HTMCs. **(A)** Primary human trabecular meshwork cells (HTMCs, n=3) were pretreted with PBA, NaB, or NaAc followed by ZIKV (Z) (PRVABC59 strain, MOI: 1) infection for 48h. Mock-treated cells were used as controls. Infected and mock-treated cells were fixed and immunostained for anti-flavivirus E (4G2) antigen. The representative images show the presence of ZIKV (Red) and DAPI-stained nucleus (Blue), scale bar: 50µm. **(B)** In a second set of experiments, cell lysates from ZIKV-infected, PBA, NaB, NaAc, and mock-treated cells were subjected to immunoblotting for ZIKV nonstructural protein, NS3. **(C)** Densitometric analysis of NS3 immunoblots was performed using ImageJ. The bar graph represents the mean ± SD of three biological replicates. **(D)** The treated and untreated conditioned media were subjected to plaque assay to quantify the replicating virions. The bar graph represents mean ± SD from three biological replicates. ****, P < 0.0001 (one-way ANOVA).

### SCFAs butyrate and acetate diminish ZIKV-induced pro-inflammatory cytokines and IFNs/ISGs response

SCFAs have been demonstrated to modulate inflammatory and antiviral responses, although different studies have shown contentious results. SCFA butyrate has been shown to suppress inflammatory cytokines IL-1β, IL-6, and TNFα in HSV1(32–34), and Japanese encephalitis virus(36) infection; IFNs/ISGs in IA virus infection(30), whereas acetate treatment has been shown to increase IFN-β production upon RSV infection(37). However, how SCFAs modulate inflammatory and antiviral responses upon ZIKV infection remains unknown. Previously, we demonstrated that ZIKV infection in HTMC induces a dysregulated cytokine and IFN response(11). Thus, to gain molecular insight into the effect of butyrate and acetate on the inflammatory cytokine and IFNs/ISGs response to ZIKV infection, we challenged HTMC cells with and without PBA/NaB/NaAc and measured the host innate inflammatory and antiviral response 48h post-infection via qPCR. Our results show that NaAc, PBA, and NaB significantly reduced the mRNA expression levels for classical pattern recognition receptors (PRRs) (e.g., RIG-I, TLR3, and MDA5), inflammatory cytokines/chemokines (e.g., IL-6, IL-1β, and CCL-4), IFNs (e.g., IFN-α2, IFN-β1, and IFNγ), and ISGs (ISG15, OAS2, and MX1) as compared to ZIKV-infected cells (**Fig. 2**).

**Figure 2:**
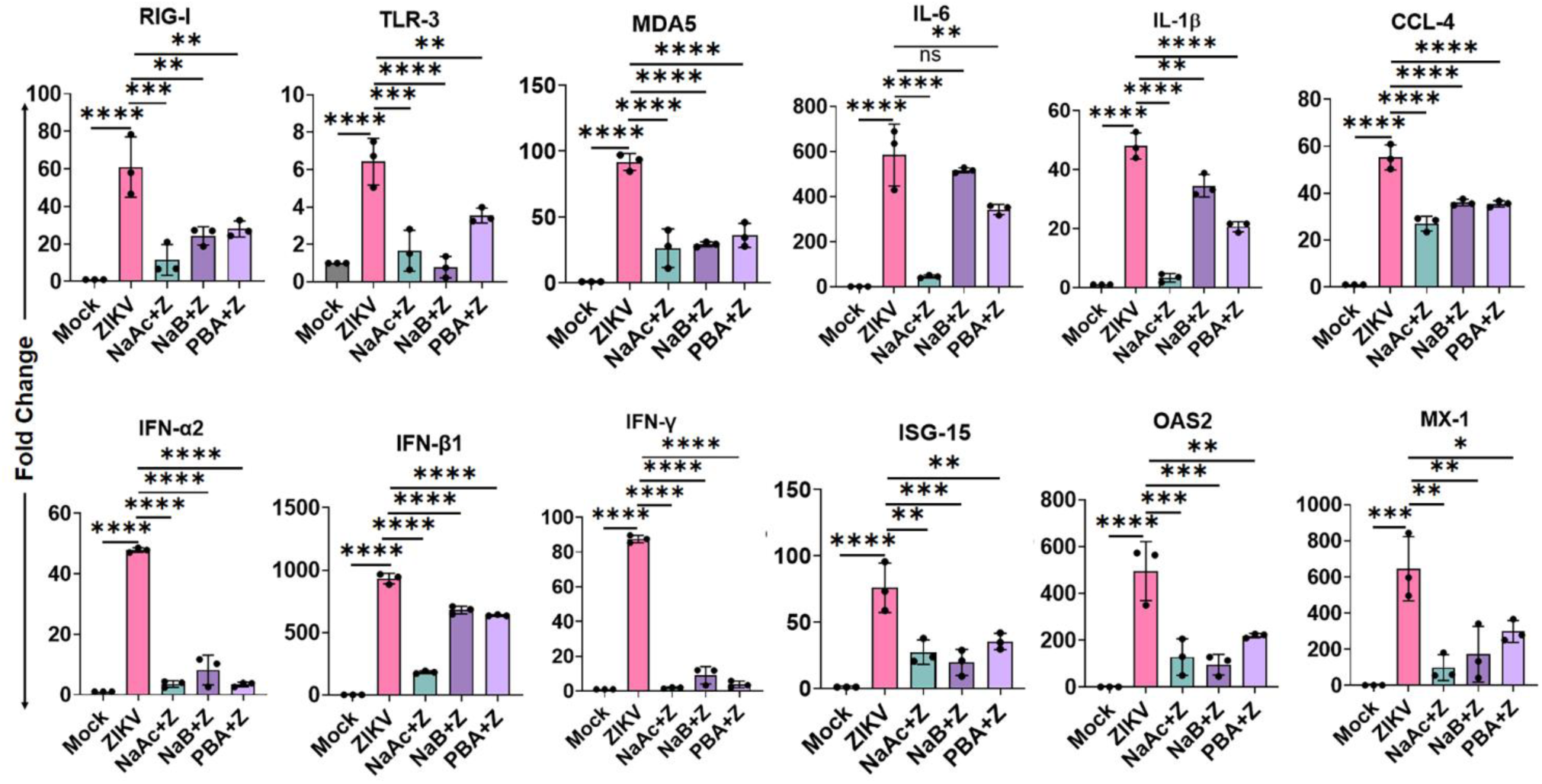
SCFA treatment attenuates ZIKV-induced pro-inflammatory cytokines and IFNs/ISGs response. HTMCs (n=3) were pretreted with PBA, NaB, or NaAc followed by ZIKV (Z) (PRVABC59 strain, MOI: 1) infection for 48h. Mock-treated cells were used as controls. Total RNA extracted from infected and treated cells were subjected to qPCR to quantify the mRNA expression of PRRs (RIG-I, TLR3, MDA5), inflammatory cytokines/chemokines (IL-6, IL-1β, CCL4), IFNs (IFN-α2, IFN-β1, IFNγ), and ISGs (ISG-15, OAS2, MX-1) genes. The bar graph represents means ± SD from three biological replicates. * P < 0.05; ** P < 0.005; *** P < 0.0005; ****, P < 0.0001 (one-way ANOVA).

NFκB, MAPK, and STAT signaling regulate the expression of viral-mediated cytokines, IFNs, and ISGs responses. Thus, to identify the mechanism by which NaAc, PBA, and NaB block the expression of these PRRs, cytokines, IFNs, and ISGs, we measured the activation of NFκB, MAPK (ERK1/2 and p38), STAT1, STAT2, and STAT3 signaling pathways in the presence and absence of these SCFAs following ZIKV infection. The western blot analysis of ZIKV-infected HTMC demonstrated increased phosphorylation of NFκB, ERK1/2, p38, STAT1, STAT2, and STAT3, (**Fig. 3**). In contrast, NaAc, NaB, and PBA treatment, significantly inhibited the activation of all of these signaling pathways (**Fig. 3**). We next assessed whether butyrate and acetate regulated the expression of antiviral/ISGs mediators and found that PBA, NaB and NaAc treatment suppressed the expression of critical antiviral/ISGs mediators, such as RIG-I, IRF3, and IFIT2 (**Fig. 3**). Together, our data indicate that SCFAs butyrate and acetate inhibits the expression of pro-inflammatory cytokines, interferons, and ISGs by inhibiting the NFκB, MAPK, and STAT1/2/3 signaling pathways.

**Figure 3:**
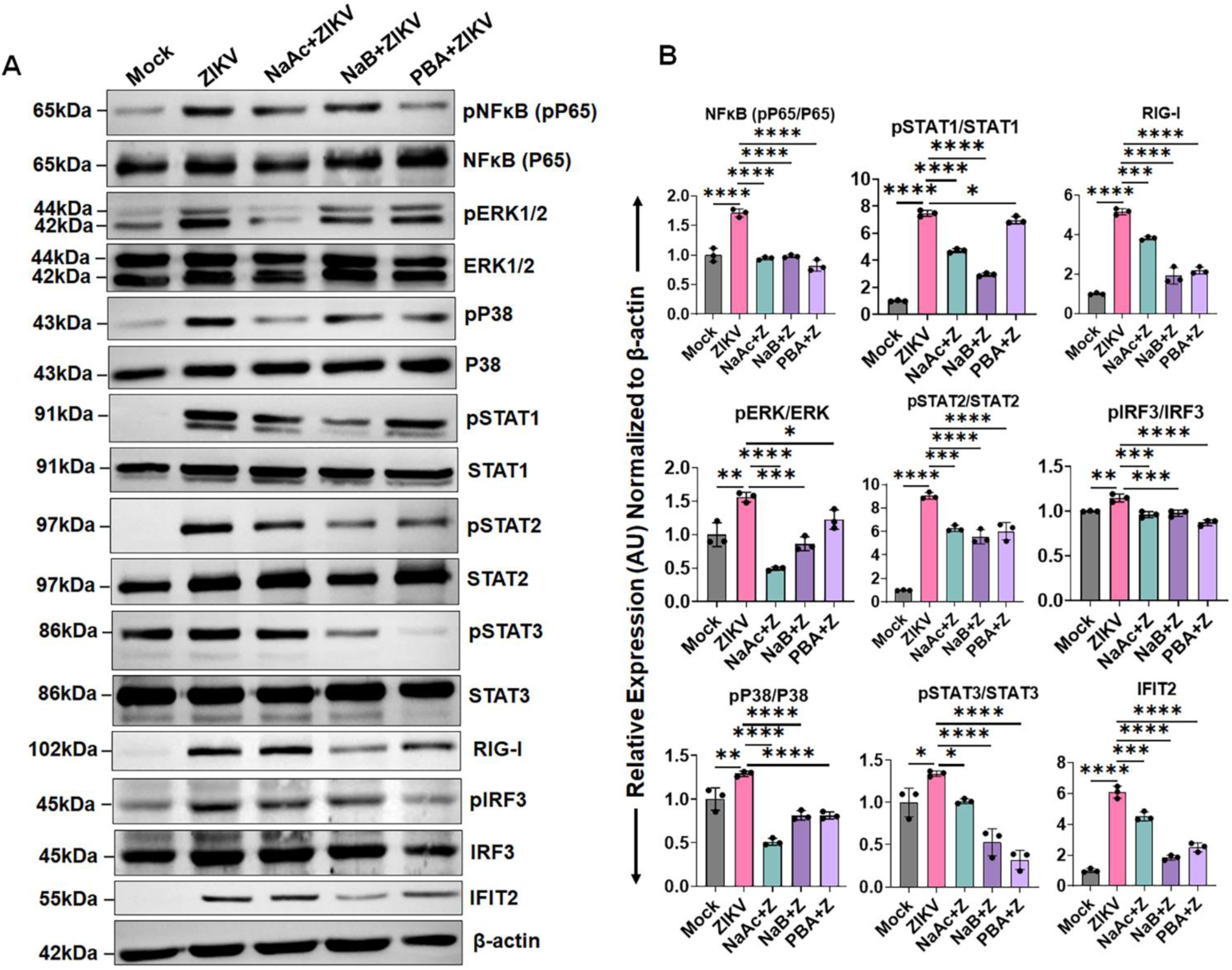
SCFAs antagonize NF-κB/MAPK-mediated ISGs signaling. HTMCs (n=3) were pretreted with PBA, NaB, or NaAc followed by ZIKV (Z) (PRVABC59 strain, MOI: 1) infection for 48h. Mock-treated cells were used as controls. The cell lysates from mock, ZIKV-infected, and drug-treated cells were subjected to western blotting for the NFκB, ERK1/2, p38, STAT1, STAT2, STST3, RIG-I, IRF3, and IFIT2 pathways. **(B)** Densitometric analysis of the immunoblots was performed using ImageJ. The bar graph represents means ± SD from three biological replicates. * P < 0.05; ** P < 0.005; *** P < 0.0005; ****, P < 0.0001 (one-way ANOVA).

### FFAR2/GPCR43 inhibition enhances the HTMC susceptibility towards ZIKV

SCFAs, acetate and butyrate, are known ligands for FFAR2/GPR43. FFAR2 is expressed by the intestinal epithelium, brain, and various subsets of immune cells such as T cells, monocytes, macrophages, neutrophils, and dendritic cells(31, 38). However, their expression by the trabecular meshwork (TM) is unknown. Activation of FFAR2 by SCFAs has been shown to diminish susceptibility toward various microbial pathogens, including bacteria and viruses(37, 39), but their role in flaviviral infections such as ZIKV has never been demonstrated. Therefore, here we aimed to investigate the role of FFAR2 in ZIKV infectivity. To assess the role of FFAR2 in HTMC, we pretreated the cells with PBA and NaAc prior to ZIKV infection and measured the expression of FFAR2 via qPCR, western blotting, and immunofluorescence staining. Our results from all three assays demonstrated that ZIKV significantly induces the expression of FFAR2 at transcript as well as protein levels in HTMC, which is potentiated by its ligands PBA and NaAc (**Fig. 4A, B & C**). To further test the role of FFAR2 in ZIKV infectivity, we blocked the FFAR2 receptor using a selective pharmacological inhibitor of FFAR2, 2-(4-chlorophenyl)-3-methyl-*N*-(thiazol-2-yl)butanamide (4-CMTB)(31, 40). Our data shows that inhibition of FFAR2 substantially increased the ZIKV replication in HTMC, even in the presence of PBA/NaAc, as revealed by increased ZIKV-4G2 immunofluorescence staining (**Fig. 4D**) and ZIKV-NS3 immunoblotting (**Fig. 4B**). The inhibition of the FFAR2 receptor by 4-CMTB was confirmed by qPCR, western blotting, and immunostaining (**Fig. 4A, B & C**). Together, our results indicate that PBA and NaAc mediate their antiviral activity via FFAR2 signaling, and inhibition of FFAR2 dramatically enhances the HTMC susceptibility towards ZIKV infection.

**Figure 4:**
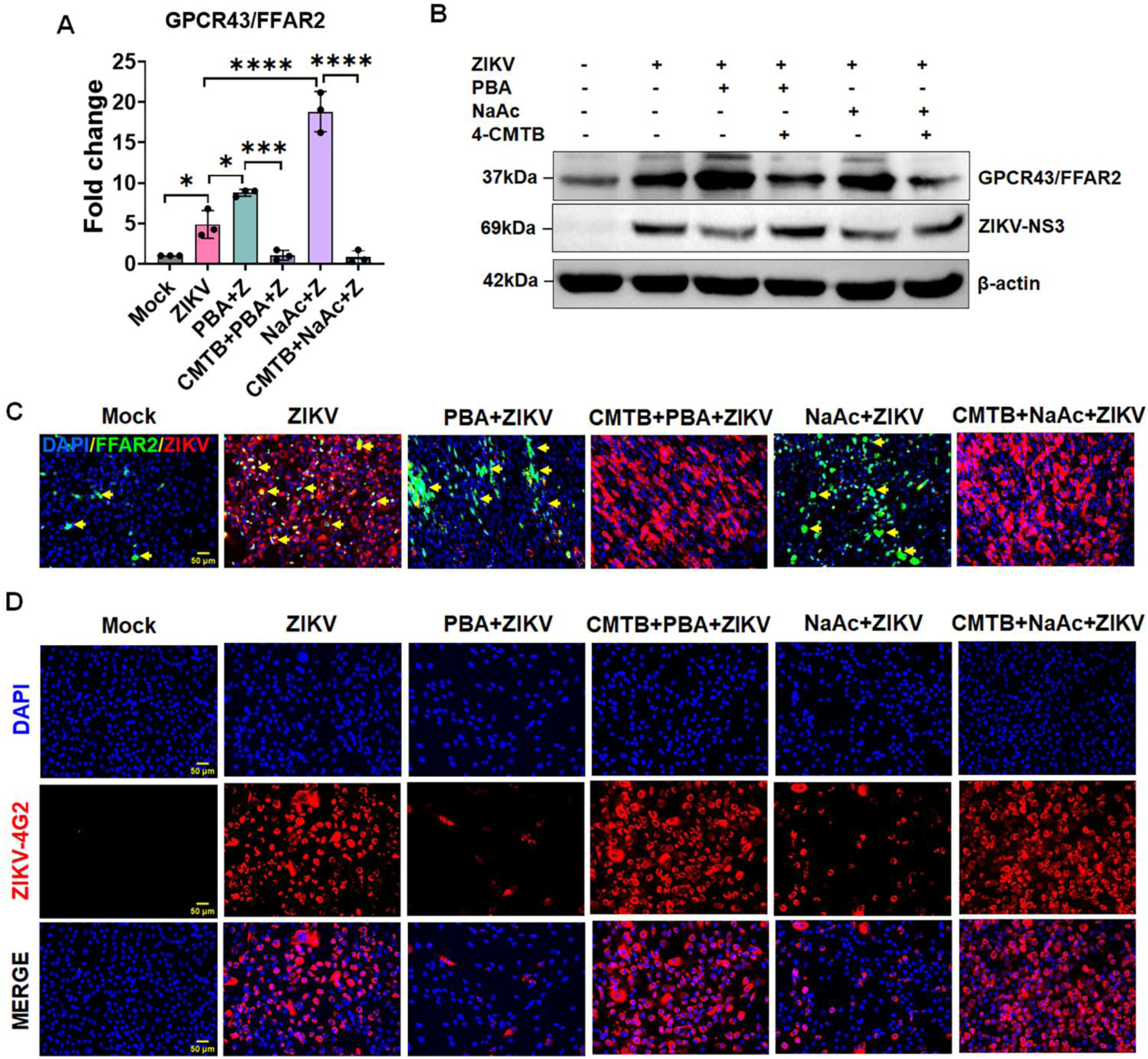
HTMC expresses FFAR2, and its inhibition promotes ZIKV replication. HTMCs (n=3) were pretreated (1h) with FFAR2 inhibitor 4-CMTB, followed by PBA, NaB, or NaAc treatment for another 1h. After pre-treatment, cells were infected with ZIKV (Z) (PRVABC59 strain, MOI: 1) for 48h. Mock-treated cells without infection were used as controls. **(A)** Total RNA extracted from mock, ZIKV-infected, and drug-treated cells were subjected to qPCR to quantify the mRNA expression of FFAR2 gene. **(B)** From another set of experiments, the cell lysates of infected and treated cells were subjected to western blotting for the FFAR2 and ZIKV NS3 proteins. **(C)** Cells were fixed and co-immunostained for FFAR2 and ZIKV E antigen (4G2) antibodies. The representative images show staining for FFAR2 (Green), ZIKV (Red), and DAPI-stained nucleus (Blue), scale bar: 50µm. **(D)** In another set of experiments, cells were fixed and immunostained for the presence of ZIKV (Red), DAPI-stained nucleus (Blue), scale bar: 50µm.

### SCFA treatments suppress, while FFAR2 inhibition promotes ZIKV-induced cell death

Previously, we demonstrated that ZIKV induces TM cell death *in vitro* as well as in mouse eyes(11). Here, we sought to test if SCFAs PBA, NaB, and NaAc have any effect on TM cell death and if FFAR2 inhibition modulates ZIKV-induced cell death. To test this, we infected HTMC with ZIKV in the presence and absence of PBA and NaAc with and without FFAR2 inhibition and performed a TUNEL assay to assess cell death. Our results show that PBA and NaAc treatment significantly protected the HTMC from ZIKV-induced cell death, as evidenced by decreased TUNEL-positive cells compared to ZIKV-infected/untreated cells (**Fig. 5A & B**). However, the FFAR2 blocking abolished the cell protective ability of PBA and NaAc and significantly enhanced the ZIKV-induced cell death (**Fig. 5A & B**). Our findings indicate that PBA and NaAc protect HTMC from ZIKV-induced cell death, and the cell protective ability of these SCFAs is mediated via FFAR2.

**Figure 5:**
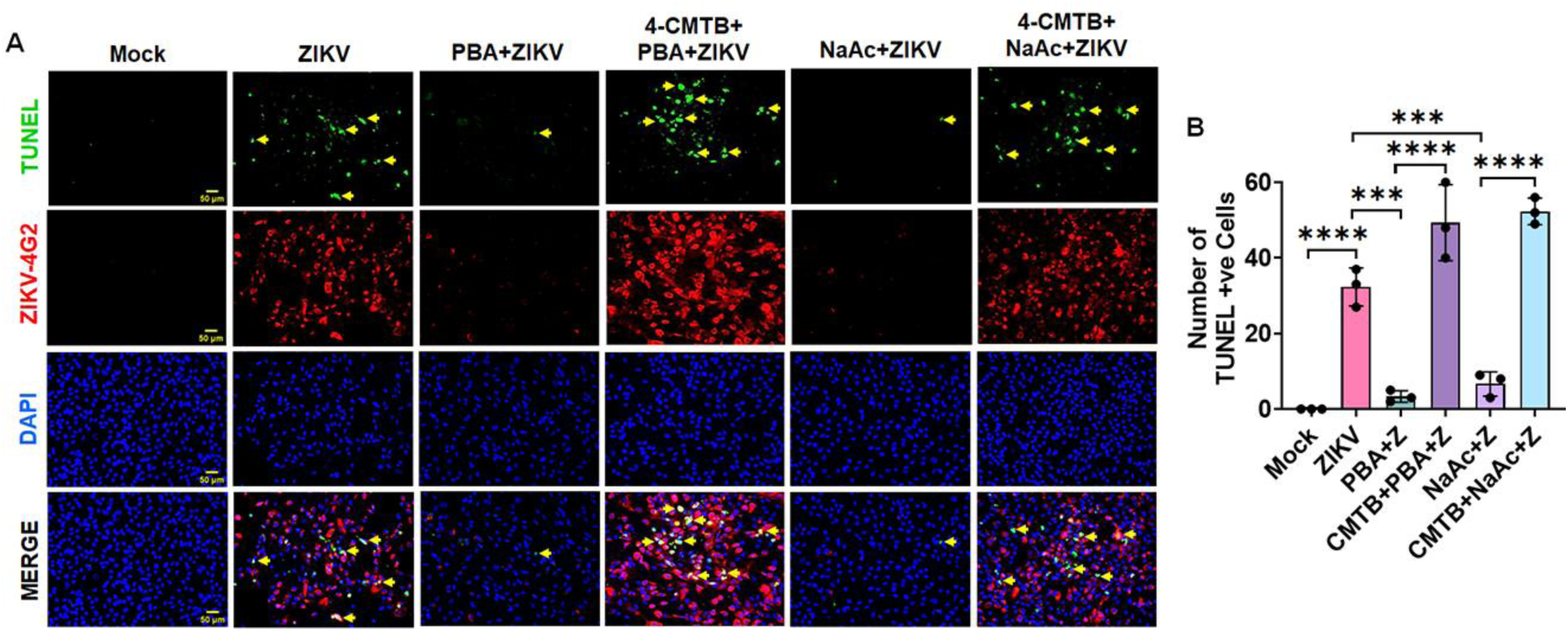
SCFA treatments suppress while FFAR2 inhibition promotes ZIKV-induced cell death. HTMCs (n=3) were pretreated (1h before) with FFAR2 inhibitor 4-CMTB, followed by PBA, NaB, or NaAc treatment for 1h. After pre-treatment, cells were infected with ZIKV (Z) (PRVABC59 strain, MOI: 1) for 48h. Mock-treated cells without infection were used as controls. **(A)** Cells were fixed and subjected to TUNEL assay followed by co-immunostaining for ZIKV E (4G2). Representative images show TUNEL-positive cells (Green-a few representative marked with yellow arrow) with ZIKV (Red) and DAPI nuclear stain (Blue). Scale bar: 50µm. **(B)** The average number of TUNEL-positive cells per field (average of 4 frames/slides from n=3) were counted and presented as mean ± SD of TUNEL-positive cells. *** P < 0.0005; ****, P < 0.0001 (one-way ANOVA).

### SCFAs exhibit anti-ZIKV activity by direct inactivation and reduced viral binding to cells

SCFAs have shown differential antiviral activities against different viruses. Some studies have shown that SCFA supports viral replication by suppressing the interferon response, while some have suggested suppressed viral replication despite the reduction of inflammatory and antiviral response(30, 41). We were puzzled by our observations that butyrate and acetate inhibit ZIKV replication in HTMC as well as suppress the inflammatory/IFNs/ISGs response. This indicates an alternative antiviral mechanism of SCFAs independent of IFN/ISG signaling. Thus, we aimed to determine the mode of action of these SCFAs on ZIKV. SCFA has been shown to influence viral entry, replication, and reactivation(31, 42, 43). Therefore, we decided to test if SCFAs butyrate and acetate alter the ZIKV binding to HTMC and inactivate the virus. To test this, we performed viral attachment and entry assays by measuring the viral RNA copy numbers and direct inactivation via plaque assay following incubation with SCFAs as described in the methodology section. Our results from the viral attachment assay revealed a significant decrease in viral copy numbers with SCFA treatment (**Fig. 6A**). Similarly, the entry assay also demonstrated a substantial decline in viral copy numbers (**Fig. 6B**). To test the viral inactivation by SCFA, we incubated ZIKV with PBA, NaB, and NaAc for 2h and 4h in culture media without cells and estimated the number of replicating viruses via plaque assay. Our data revealed that incubation of ZIKV with these SCFAs significantly reduced the number of plaques (>two log_10_ fold), indicating viral inactivation (**Fig. 6C**).

**Figure 6:**
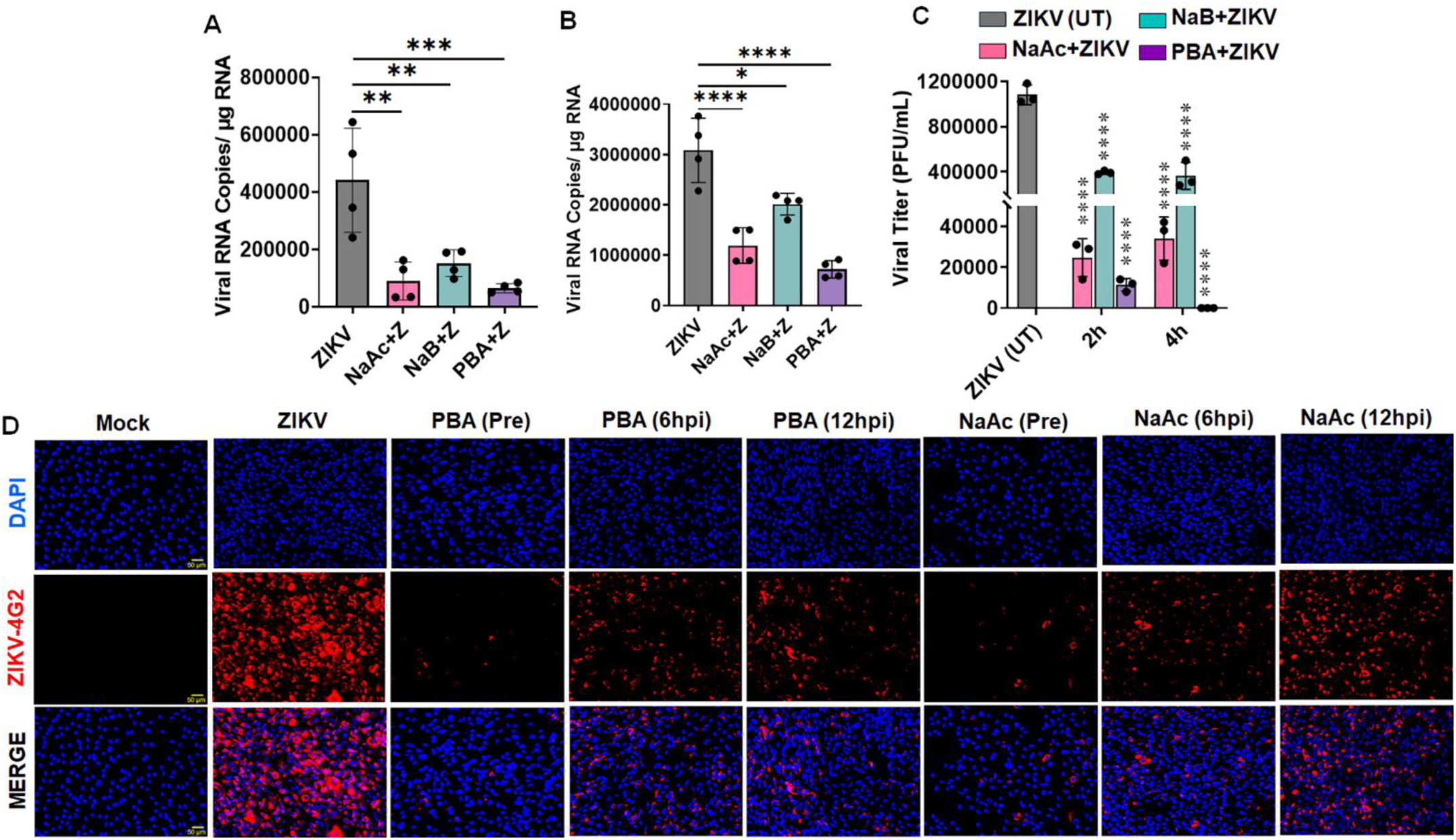
SCFA inhibits ZIKV infection by viral inactivation and inhibition of viral attachment to the cells. **(A)** HTMC (n=4) were treated with PBA, NaB, and NaAc for 1 h with incubation at 4°C, followed by ZIKV (MOI: 1) infection. Following incubation at 4°C for 2h, the total RNA was extracted and subjected to qPCR for the ZIKV envelope gene. The data were presented as mean ± SD of RNA copies/μg RNA. **(B)** HTMC (n=4) were treated with PBA, NaB, and NaAc for 1 h with incubation at 37°C, followed by ZIKV (MOI: 1) infection. After viral adsorption for 2h at 37°C, the total RNA was extracted and subjected to qPCR for the ZIKV envelope gene. The data were presented as mean ± SD of RNA copies/μg RNA. **(C)** PBA, NaB, or NaAc were incubated with ZIKV (10^6^ PFU/mL) in a serum-free DMEM for 2h and 4h at 37°C. ZIKV without drugs was used as an untreated (UT) control. At respective timepoints, the media were collected and subjected to plaque assay for viral titer estimation. The plaques from treated and untreated conditioned media were counted and presented as PFU/mL. **(D)** HTMC were either pretreated (1h before) with PBA or NaAc, followed by ZIKV infection, or treated 6h and 12h post-ZIKV infection. Mock-treated and ZIKV-infected cells without treatment were used as controls. Forty-eight hours after infection, cells were fixed and immunostained for ZIKV E (4G2) antigen. The representative images show the presence of ZIKV (Red) and DAPI-stained nucleus (Blue), scale bar: 50µm. * P < 0.05; ** P < 0.005; *** P < 0.0005; ****, P < 0.0001 (one-way ANOVA).

To further confirm the viral inactivation before entry by SCFAs, we tested the anti-ZIKV activity of PBA and NaAc in pre-(before viral internalization) and 6 and 12h post-infection (after viral internalization) treatment settings. We observed that PBA and NaAc suppressed the viral replication in pretreated groups; however, ZIKV positivity was considerably higher in post-treatment settings in comparison to the pretreated groups (**Fig. 6D**). Though both pre- and post-treatment with butyrate and acetate suppressed the viral replication up to some extent compared to ZIKV-infected and untreated cells (**Fig. 6D**). These results indicate that the anti-ZIKV activity of SCFA could be partially due to the inhibition of viral binding and inactivation before internalization.

### SCFA treatment attenuates while FFAR2 inhibition exacerbates ZIKV-induced ocular manifestations in mice

Previously, we showed that ZIKV induces anterior segment (AS) inflammation leading to TM damage and elevated intraocular pressure(11). We also demonstrated that ZIKV migrates from the anterior chamber to the back of the eye and causes chorioretinal atrophy, retinal and optic nerve damage(10, 11). After observing the anti-ZIKV effect *in vitro* in HTMC, we reasoned to test their therapeutic potential against ZIKV-induced ocular complications in mice. To test this, we pretreated (one day before) mice with PBA and NaAc via i.p. administration, followed by ZIKV infection and subsequent treatment for four consecutive days. For FFAR2 inhibition groups, we pretreated mice with 4-CMTB via i.p. injections one day before PBA/NaAc pre-treatment and once again on the third day after the first administration for consistent FFAR2 blocking. The timeline for the 4-CMTB/SCFAs treatment and ZIKV infection is shown in **Fig. 7A**. As anticipated, ZIKV infection significantly elevated the IOP (**Fig. 7B**), caused severe RPE/chorioretinal atrophy (**Fig. 7C**), and RPE/outer retinal layer disruption (**Fig. 7D**) in comparison to uninfected controls. In contrast, PBA and NaAc treatment significantly reduced the IOP (**Fig. 7B**), and attenuated the ZIKV-induced RPE/retinal damage as revealed by fundus (**Fig. 7C**) and OCT (**Fig. 7D**) imaging. Remarkably, pharmacological inhibition of FFAR2 by 4-CMTB antagonizes the protective ability of PBA/NaAc and aggravates ZIKV-induced RPE/chorioretinal atrophy, outer retinal layer disruption, and IOP elevation (**Fig. 7B-D**). To further test the role of PBA and NaAc and their receptor FFAR2 on ZIKV-induced AS inflammation, we measured the expression of various inflammatory mediators in the AS tissue via qPCR with and without FFAR2 inhibition. Our result reveals that butyrate and acetate treatment significantly diminished the mRNA expression of ZIKV-induced PRRs (RIG-I, TLR3), inflammatory cytokines/chemokines (IL-6, IL-1β, CCL4), IFNs (IFN-α2, IFN-β1), and ISGs (ISG15, OAS2, MX1) in mouse AS tissue (**Fig. 8A**). Blocking FFAR2 receptor by 4-CMTB reversed the anti-inflammatory effect of PBA and NaAc, and significantly enhanced the expression of these inflammatory mediators. Since we observed an antagonizing effect on inflammatory and ISGs pathways by PBA and NaAc *in vitro*, we tested the role of these SCFAs on these pathways in our *in vivo* model. Similar to HTMCs, PBA/NaAc inhibited the NFκB, MAPKs (p38, ERK1/2), and STATs (STAT1, STAT3) and ISGs (RIG-I, IRF7) pathways activation in mouse AS/TM tissue, while FFAR2 inhibition reversed these inhibitory properties **(Fig. 8 B&C)**. Collectively, these findings indicate that SCFA suppresses ZIKV-induced AS inflammation and ocular manifestations, and the *in vivo* protective effect of acetate and butyrate is also mediated via FFAR2.

**Figure 7:**
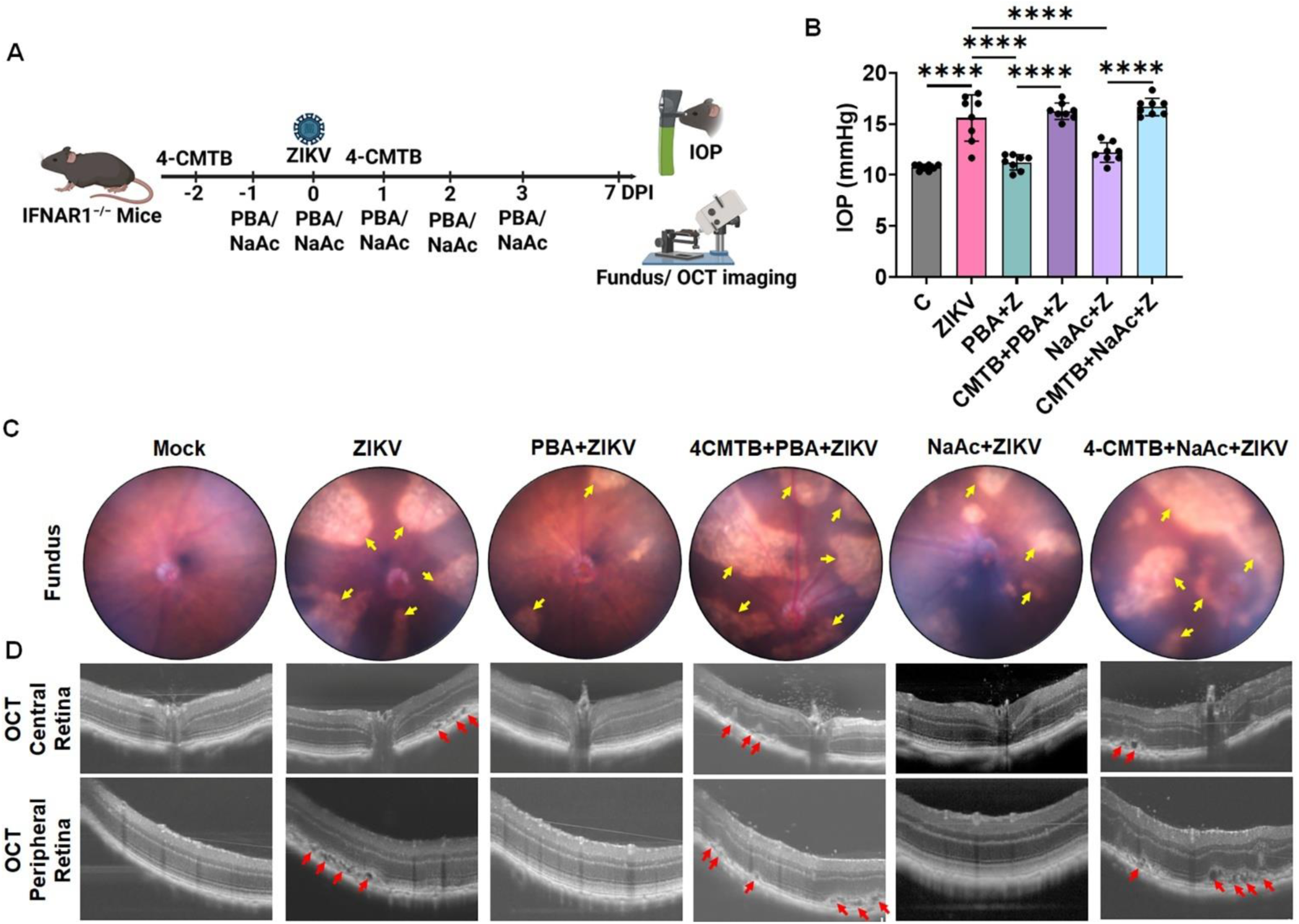
SCFA treatment attenuates while FFAR2 inhibition aggravates ZIKV-induced ocular pathology in mice. IFNAR1^−/−^ mice (n=6-8) were pretreated with FFAR2 inhibitor 4-CMTB (days −2 and 1), followed by PBA, NaB, or NaAc [days −1 to 3 days post infection (DPI)] via i.p. administration. Following treatment, mice were infected with ZIKV (1×10^4^ PFU) via intracameral injection. Mice with saline injection without ZIKV infection were used as mock controls. **(A)** Timeline for drug administration and ZIKV infection. **(B)** At 7DPI, mouse IOP were recorded and represented as mean IOP ± SD. **(C)** Fundus and **(D)** OCT imaging were performed to determine the retinal manifestations using the Micron IV fundus camera and image-guided OCT2 system. Representative funduscopic and OCT micrographs show RPE and chorioretinal atrophy [pointed with Yellow arrows (on fundus images, **C**)], with RPE and outer retinal layer disruption [pointed with Red arrows (on OCT images, **D**) in treated/ untreated groups. The timeline shown in panel A was created using BioRender software (<u>biorender.com</u>)

**Figure 8:**
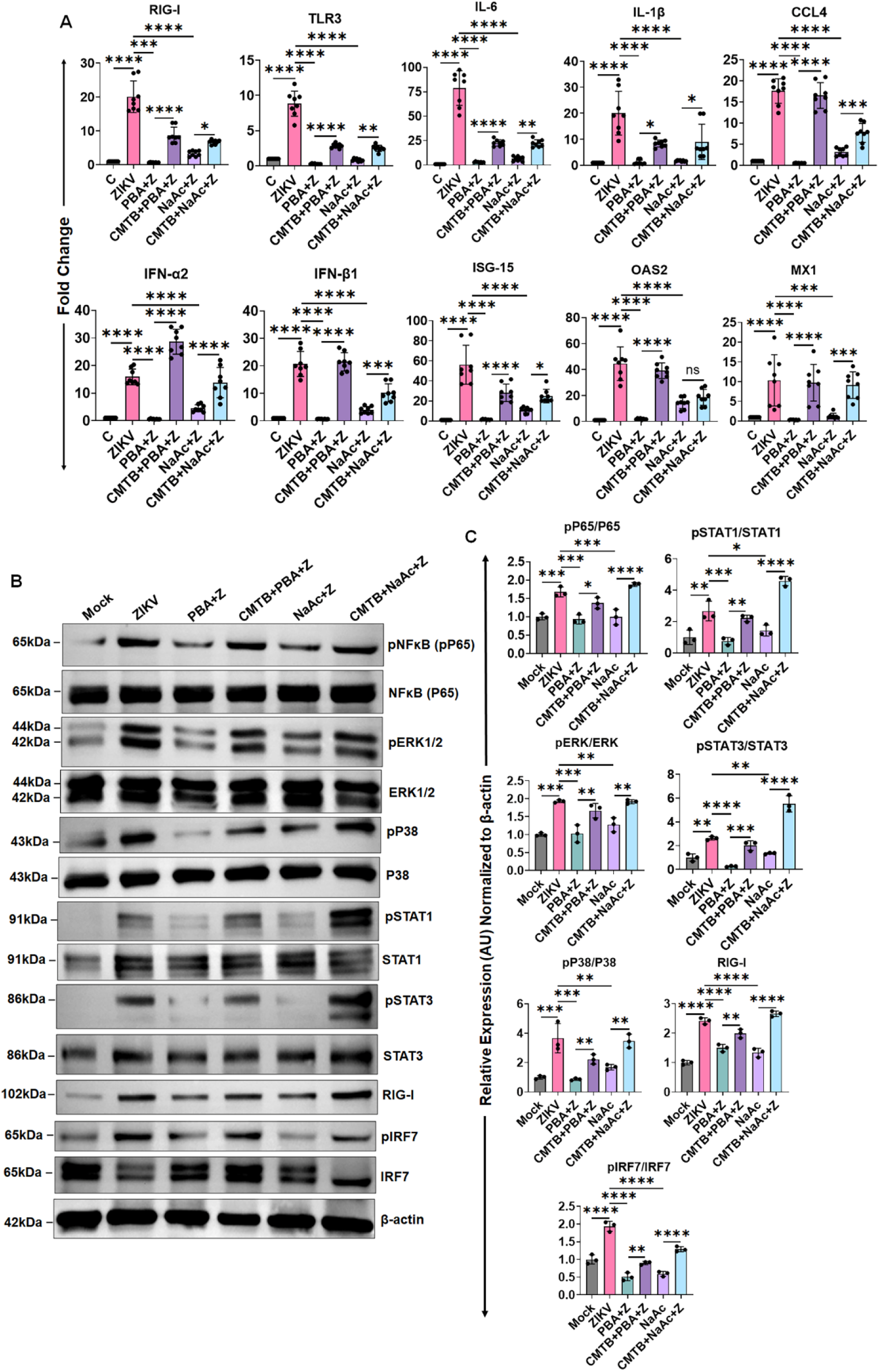
SCFA treatment diminishes while FFAR2 inhibition exacerbates ZIKV-induced mouse anterior segment’s inflammatory response. IFNAR1^−/−^ mice (n=6-8) were pretreated with FFAR2 inhibitor 4-CMTB followed by PBA, NaB, or NaAc via i.p. administration, and ZIKV infection as per the timeline shown in Figure 7A. Seven days post-infection, anterior segment/TM tissue from treated and untreated mice were harvested and subjected to **(A)** RNA extraction and qPCR to measure the mRNA expression levels of PRRs (RIG-I, TLR3), inflammatory (IL-6, IL-1β, CCL-4), IFNs (IFN-α2, IFN-β1), and ISGs (ISG15, OAS2, MX1), and **(B)** western blot for NFκB, MAPKs (ERK1/2, p38), STAT1, STAT3, RIG-I, and IRF7 inflammatory/ISG pathways. **(C)** Densitometric analysis of the immunoblots was performed using ImageJ. The bar graph represents means ± SD from three biological replicates. * P < 0.05; ** P < 0.005; *** P < 0.0005; ****, P < 0.0001 (one-way ANOVA).

## Discussion

Recent advancements in the microbiome-immunity axis have uncovered the involvement of gut microbiomes and their metabolites in multiple diseases. A recent study demonstrated alterations in gut microbiota in immunocompetent mice(44), and enhanced peripheral and nervous system inflammation due to decreased SCFA levels in macaques(45) during ZIKV infection. SCFAs, acetate, butyrate, and propionate have demonstrated promising therapeutic potential against multiple microbial pathogens, including bacteria and viruses. Although due to both pro- and antiviral properties, SCFAs have shown very complex and contrary roles with different viruses(17). Emerging literature suggests an interplay between gut microbiota and ocular health(46, 47). However, the role of SCFAs against ZIKV ocular infection remains unknown. Here, we showed a previously unreported role of butyrate and acetate and their receptor FFAR2 in ZIKV ocular pathogenesis. In this study, we discovered that SCFAs, PBA, NaB, and NaAc restrict ZIKV transmission and modulate viral attachment and internalization to the cells. We further uncovered that PBA and NaAc mediate their anti-ZIKV activity via FFAR2, and pharmacological inhibition of FFAR2 antagonizes the protective abilities of PBA and NaAc. SCFA treatment attenuates ZIKV-induced pro-inflammatory response, cell death, and ocular complications.

ZIKV infection is linked to multiple congenital malformations, including microcephaly and ocular manifestations. *In utero* exposure to ZIKV caused severe ocular abnormalities, including chorioretinal atrophy, RPE mottling, optic nerve damage, and congenital glaucoma(10, 14–16). Trabecular meshwork (TM) regulates the intraocular pressure (IOP), and pathologic stress may cause TM damage, resulting in increased glaucoma phenotype(11, 48). Recently, we discovered that ZIKV has high tropism towards the anterior segment of the eye and can cause TM damage, resulting in increased IOP and retinal ganglion cell (RGC) loss(11). In this study, we aimed to test whether SCFAs can protect TM from ZIKV infection. We found that SCFA derivatives PBA, NaB, and NaAc dramatically reduced ZIKV replication and transmission. We further discovered that butyrate and acetate inhibit viral binding and cellular entry to the TM cells. The SCFAs also protected TM from ZIKV-induced cell death. Our study corroborated with previous findings where SCFA treatment has been shown to suppress replication of SARS-CoV-2(22), HBV(35), HSV-1(32–34), and porcine epidemic diarrhea virus (PEDV)(49), and exacerbate disease severity. Similarly, butyrate has been shown to effectively reduce rotavirus-induced cell death and provide protection against intestinal epithelial barrier damage(43, 50). In contrast to these findings, a few other studies have shown that butyrate could promote the replication of EBV(51), IAV, HIV-1, hMPV, VSV(30), and TGEV(52), whereas it has no effect on SeV replication(30). These findings suggest that, given the diverse role of SCFA among different viruses, it is imperative to interpret their pro- or anti-viral activity within specific relevant physiological contexts.

Upon viral infection, host cells induce a pro-inflammatory and IFN response, which plays a crucial role in curbing infection. However, uncontrolled immune activation upon infection may lead to persistent inflammation, resulting in tissue damage, morbidity, and mortality. The recurrent inflammation of ocular tissue, an immune-privileged organ, is detrimental and results in vision loss. ZIKV has been shown to induce inflammatory cytokines, IFNs, and ISGs in different models(53). We recently observed a dysregulated immune response via ZIKV in TM, resulting in trabeculitis and TM damage(11). SCFAs are well-characterized for their anti-inflammatory activities via HDAC inhibition(17, 52). Recent studies have demonstrated that SCFAs can suppress inflammatory cytokines and virus-induced IFN/ISG response to confer protection from tissue damage(17, 33, 34). In line with these studies, we observed that PBA, NAB, and NaAc can suppress ZIKV-induced inflammatory (IL-6, IL-1β, CCL-4), IFNs (IFNα, IFNβ, IFNγ), and ISGs (ISG15, OAS2, MX1) responses in TM. During RNA virus infection, the inflammatory and antiviral immune response is initiated upon PRRs (RIG-I, MDA5, TLR3), followed by activation of NFκB/MAPK/STAT pathways. ZIKV activates RIG-I, MDA5, and TLR3 receptors and mediates innate immune response via NFκB/MAPK pathways(54, 55). SCFA treatments have been shown to antagonize the NFκB pathway upon HSV-1(34) and Japanese encephalitis virus(36) infection. In this study, we observed that PBA, NaB, and NaAc suppressed the activation of PRRs (RIG-I, TLR3, MDA5), NFκB, MAPKs (ERK1/2, p38), and IFNs/ISGs mediators, including STATs (STAT1, STAT2, STAT3), IRF3, and IFIT2. Similar to our findings, butyrate has shown inhibition of IFNβ-induced RIG-I and IFIT-2 expression(30). In contrast, this study reported that butyrate has no effect on STAT1 and STAT2 upon IFNβ stimulation. This study also concluded that butyrate has a differential role in ISG expression; it suppressed 60% of IFN-induced ISGs while upregulating 3% of IFN-induced ISGs(30). Another study by He et.al. reported increased IFN and ISG production by butyrate against PEDV in porcine intestinal epithelial cells(49). Similarly, acetate has shown enhanced IFNβ production against RSV(37) and NLRP3-IFN-I mediated antiviral response against IAV(56). However, our finding corroborated with other studies showing suppression of JAK/STAT pathways in LPS-stimulated corneal fibroblasts(57) and RIG-I signaling in TGEV and SARS-CoV-2 infection models(52, 58). These differential effects could be potentially due to either the stressor (IFNβ vs PEDV vs ZIKV infection) or the cell models (transformed colon/lung cell line vs porcine epithelial cells vs human primary TM) used in these studies, versus our study. As discussed by Yin et.al, the differential effect of SCFA could also be attributed to the distinct SCFA treatment methods(52). Indeed, we observed alteration of viral binding and cellular entry by SCFAs, and therefore differential effects in viral dissemination in pre-vs. post-treatment. Similarly, one other study has demonstrated the differential impact of acetate treatment in mice with simultaneous, before, or after IAV infection(56). Collectively, these findings imply the pleiotropic role of SCFAs on viral infection and immune function and underline careful consideration when applying the findings to human patients with different viral infections.

Canonically, SCFAs mediate their activity by activating the GPCRs, also known as the free fatty acid receptors (FFAR). FFAR2 is expressed by multiple cells, including immune cells, skeletal muscles, smooth muscles, and neurons, and therefore assists in maintaining homeostasis by SCFAs in the colon, kidney, joints, lungs, and brain. In the eye, FFAR2 is expressed by corneal epithelium and endothelium, iris, ciliary body, inner nuclear, outer nuclear, and ganglion cell layers(18). FFAR2 plays a critical role in regulating gut homeostasis, energy utilization, immune cell functions, and suppression of inflammatory response. It has also been shown to play a protective role in non-infectious ocular diseases such as uveitis(18). However, their role in ZIKV ocular pathogenesis is entirely unknown. Here, we observed for the first time that TM cells express the FFAR2 receptor and ZIKV modulates its expression. Our study revealed that butyrate and acetate potentiated the FFAR2 expression on TM, and pharmacological inhibition of FFAR2 promoted ZIKV replication. Furthermore, FFAR2 inhibition enhanced the ZIKV-induced cell death, which was suppressed by PBA and NaAc. Our findings corroborated with previous studies where FFAR2 knockdown has shown enhanced susceptibility to *C. rodentium*(*59*), *Klebsiella pneumoniae* lung infection(60), and TLR-induced keratitis(18). Activation of FFAR2 in pulmonary epithelial cells has shown reduced virus-induced cytotoxicity(37). Similarly, targeted activation of FFAR2 has shown diminished susceptibility towards *S. aureus*, RSV, and IAV and *Streptococcus pneumoniae* superinfection(28, 37, 39, 61).

In summary, our work reveals an antiviral role of SCFAs, butyrate, and acetate, restricting ZIKV transmission and associated ocular manifestations via FFAR2 **(Fig. 9)**. The inhibition of FFAR2 reverts the protective effect of butyrate and acetate and promotes ZIKV-induced cell death. Butyrate and acetate also inhibit viral binding, cellular entry, and inactivate the virus prior to internalization. Our study could aid the future design of therapeutic interventions to treat or prevent ZIKV infection and associated ocular complications in humans. However, given the diverse role of SCFAs on different viruses, their safety must be ensured prior to therapeutic use in human subjects with various viral infections. Although we demonstrated the role of FFAR2 in ZIKV transmission using a potent and selective inhibitor of FFAR2, 4-CMTB, future studies using FFAR2 knockout are essential to decipher the role of FFAR2 in ZIKV pathogenesis. Similarly, dietary manipulations or appropriate supplements to augment SCFA production could be explored as potential antiviral strategies against ocular viral infection.

**Figure 9:**
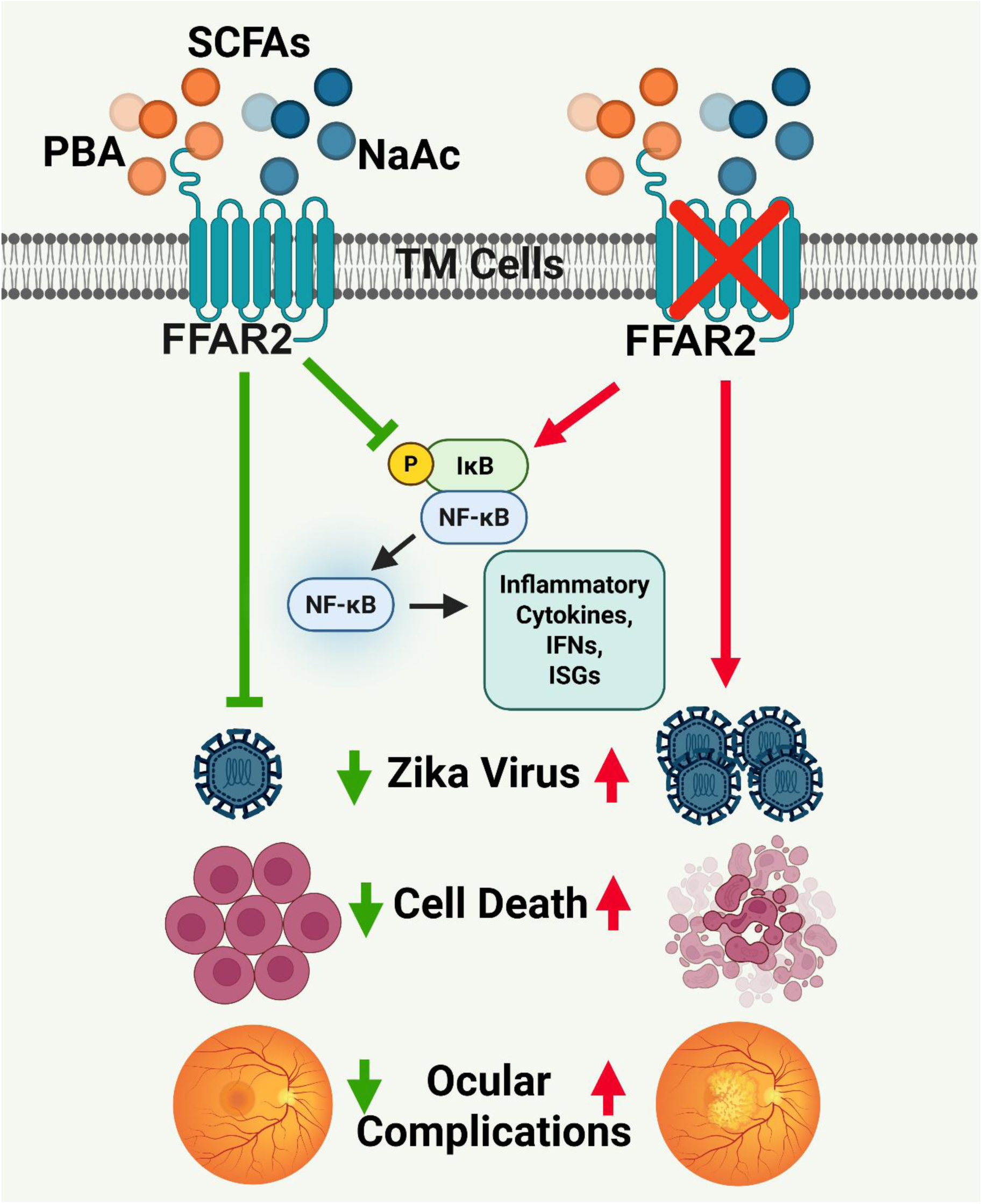
Anti-ZIKV properties of SCFA butyrate and acetate mediated via FFAR2. Butyrate and acetate suppress ZIKV transmission and associated ocular manifestations. The anti-ZIKV activity of SCFAs is mediated via FFAR2. FFAR2 inhibition promotes ZIKV replication, cell death, and ZIKV-induced ocular manifestations. The schematic diagram was created using BioRender software(<u>biorender.com</u>)

## Materials and Methods

### Antibodies and Reagents

Antibodies used in this study were purchased from the following sources: 4G2 (GeneTex, #GTX57154), NS3 (GeneTex, #GTX133309), β-actin (Millipore Sigma, #A2228), pNFκB (#3033), NFκB (#6956) pERK1/2 (#4370), ERK (#4695), pP38 (#4511), P38 (#8690), pSTAT1 (#9167), STAT1 (#14994), pSTAT2 (#88410), STAT2 (#72604), pSTAT3 (#9145), STAT3 (#9139), RIG-I (#3743), pIRF3 (#79945), IRF3 (#4302), and IFIT2 (#92633) antibodies were purchased from Cell Signaling Technology (Danvers, MA). SCFAs NaAc (Thermofisher, #A13184-30), 4-PBA (#11323), NaB (#13121), and FFAR2 receptor antagonist 4-CMTB (#29680) were purchased from Cayman Chemicals (Ann Arbor, MI).

### Cells Culture

Primary human trabecular meshwork cells (HTMC)(11) were cultured in Dulbecco’s minimal essential medium (DMEM) low glucose GlutaMAX medium, supplemented with 10% fetal bovine serum (FBS), 1X penicillin-streptomycin solution in a CO_2_ (5%) incubator at 37°C. Vero E6 cells (ATCC CRL-1586) were cultured in DMEM low glucose GlutaMAX medium containing 1X penicillin-streptomycin supplemented with 10% FBS. For virus propagation, Aedes albopictus, C6/36 cells (ATCC CRL-1660) were cultured in Eagle’s Minimum Essential Medium (EMEM) containing 1X penicillin-streptomycin supplemented with 10% FBS in a CO_2_ incubator at 30°C.

### Zika Virus Strain and Infection Procedure

The ZIKV strain PRVABC59 (NR-50240) was obtained from BEI Resources, National Institute of Allergy and Infectious Diseases (NIAID), and propagated in Aedes albopictus, C6/36 cells. The viral titers were determined using the plaque assay. Small aliquots were prepared and stored in a −80°C freezer for infection studies.

For *in vitro* experiments, HTMCs were challenged with ZIKV at a multiplicity of infection (MOI) of 1. For pharmacological inhibition/activation studies, cells were pretreated with respective drugs/antagonists for 1h, followed by virus adsorption in serum-free media for 1h, until indicated otherwise. After adsorption, cells were replenished with media containing drugs and incubated until the desired endpoint.

### Mice and Ethics Statement

IFNAR1^−/−^ mice (B6 background, MMRRC Strain # 032045-JAX) were originally purchased from Jackson Laboratories and bred in-house in a germ-free University of Missouri (MU) Office of Animal Resources (OAR) facility. Mice aged 6-10 weeks (male and female) were used in this study. Animals were housed in a restricted-access AAALAC-accredited animal facility. All animals were maintained in a 12:12 h light/ dark cycle, with free access to food (rodent chow, Labdiet Pico Laboratory, Saint Louis, MO) and water. Mice were treated in compliance with the Association for Research in Vision and Ophthalmology (ARVO) Statement for the Use of Animals in Ophthalmic and Vision Research. All biohazard and animal procedures were approved by the MU Institutional Biosafety Committee (IBC) and Animal Care and Use Committee (ACUC).

### Mouse infection and SCFA treatment

For SCFA treatment, mice were pretreated with PBA/NaAc (100 mg/kg) via i.p injections one day prior to infection and three consecutive days post-ZIKV infection. In the FFAR2 inhibition groups, mice were pretreated with 4-CMTB (10mg/kg) via i.p. injection one day prior to PBA/NaAc treatment and another dose on day three(40) since the first administration as per the timeline shown in Fig. 7A. For the ZIKV infection, anesthetized animals were inoculated with 1×10^4^ PFU of ZIKV via intracameral injections as described previously(11). Seven days post-infection, IOP was recorded using a Tonolab tonometer, and fundus imaging was performed using Micron IV (Phoenix-Micron Inc., Bend, OR). Animals were euthanized, and anterior segment tissue was harvested for inflammatory cytokines/pathways assessment via qPCR and immunoblotting.

### Plaque Assay

ZIKV plaque assay was performed using a protocol we described recently(2). Briefly, a confluent monolayer of Vero cells was infected with serial dilutions of ZIKV stock culture. One hour following viral adsorption, the cell monolayer was overlayed with the first overlay media containing a 1:1 mixture of 2X EMEM, 4% FBS, 2X P/S, 20mM MgCl_2,_ and 1.6% Noble Agar. The following day, a second overlay media containing DMEM, 1mg/ml BSA, 40mM MgCl_2_, 0.2% glucose, 2mM sodium pyruvate, 4mM L-glutamine, 4mM oxaloacetic acid, 1X P/S, and 0.1% sodium bicarbonate was added. The plates were incubated at 37°C for five days in a CO_2_ incubator. Following incubation, cells were fixed with 10% Tricarboxylic Acid (TCA) for 20 min, and the agar overlay was removed gently without disturbing the cell monolayer. Viral plaques were stained with 0.2% crystal violet for 20 min, followed by a wash with MilliQ water. The plaques were counted, and titers were estimated as log_10_ PFU/mL.

### Viral Attachment, Entry, and Inactivation Assay

The viral attachment, entry, and inactivation assay was performed as described previously(62). Briefly, HTMCs were pre-incubated with PBA (3mM)/ NaB (3mM)/ NaAc (200µM) at 4°C (for attachment assay) or 37°C (for entry assay) for one hour, followed by ZIKV infection (MOI:1). After viral adsorption for 2 h, the cells were washed three times with fresh DMEM media, and total RNA was isolated. The viral RNA copy numbers were determined from whole-cell RNA using a TaqMan probe against the ZIKV envelope (E) gene via qPCR.

For the direct inactivation assay, 1 × 10^6^ PFU/mL of ZIKV (diluted in serum-free DMEM) was incubated with PBA (3mM)/ NaB (3mM)/ NaAc (200µM) at 37°C for 2 or 4 h. ZIKV without any drug was used as an untreated control. At indicated time points, a 100µL viral-drug mixture was taken out for viral quantification via plaque assay.

### Immunofluorescence Assay (IFA)

For IFA, cells were seeded in a Nunc four-well chamber slide (Fisher Scientific, Rochester, NY) and infected with ZIKV at an MOI of 1 at ∼70-80% confluency. Mock-treated/ uninfected cells were used as controls. At desired endpoints, cells were fixed using 4% paraformaldehyde for 10 min at room temperature (RT). After three washes with 1X PBS, cells were blocked and permeabilized using 1% (w/v) BSA with 0.4% (v/v) Triton X-100 made in PBS (blocking buffer) for 1h at RT in a humidified chamber. Cells were then incubated with the primary mouse/rabbit antibodies in the blocking buffer (1:100 dilution) overnight at 4°C in a humidified chamber. Subsequently, after three washes with 1X PBS, cells were incubated with anti-mouse/rabbit Alexa Fluor 488/594-conjugated secondary antibodies (1:200 dilutions) for 1h at RT. Finally, cells were washed four times with 1X PBS and mounted in Vectashield anti-fade mounting medium containing DAPI (Vector Laboratories, Burlingame, CA). The slides were visualized and imaged using a Keyence (Keyence, Itasca, IL) fluorescence microscope.

### TUNEL Assay

The cell death was estimated using the TUNEL assay(63). HTMCs were grown in Nunc four-well chamber slides (Fisher Scientific, Rochester, NY) and infected with ZIKV in the presence or absence of SCFAs or 4-CMTB. TUNEL assay was performed using ApopTag Fluorescein In Situ Apoptosis Detection Kit (#S7110) per the manufacturer’s instructions (Millipore Sigma, Billerica, MA). Following TUNEL staining, cells were immunostained with ZIKV-4G2 antibodies using the IFA protocol described above. The cells were visualized and imaged using a Keyence microscope (Keyence, Itasca, IL).

### Immunoblotting

Immunolblotting was performed using a method we described previously(63). Briefly, the treated/untreated cells were washed using ice-cold PBS and lysed using RIPA lysis buffer containing Halt^TM^ protease and phosphatase inhibitor cocktail (Thermo Scientific, Rockford, IL). The total protein concentration was estimated using BCA protein estimation kit per the manufacturer’s instructions (Thermo Scientific, Rockford, IL). Thirty micrograms of total protein were resolved on SDS-PAGE gels and transferred onto nitrocellulose or PVDF membranes. Following transfer, the membrane was blocked using a blocking buffer containing 5% non-fat skim milk in 1X TBST. Subsequently, blots were incubated with anti-rabbit/mouse primary antibodies (1:1000 dilutions) diluted in 3% BSA overnight at 4°C with gentle agitation. After incubation with the primary antibodies, the membranes were washed three times using 1X TBST, followed by incubation with anti-mouse/rabbit HRP-conjugated secondary antibodies (1:2000 dilutions) at RT for 2h. Following three washes with 1X TBST, the blots were developed using Supersignal West Femto chemiluminescent substrate and imaged using iBright FL1500 imager (Thermo Fisher Scientific, Rockford, IL).

### RNA Isolation and Real-time qPCR

Following treatment, cells were collected in TRIzol^TM^ reagent (#15596018) and the RNA was isolated per the manufacturer’s instructions (Thermo Scientific, Rockford, IL). RNA (1µg) was reverse transcribed to cDNA using a Maxima first-strand cDNA synthesis kit, per the manufacturer’s instructions (Thermo Scientific, Rockford, IL). The cDNA was amplified using mouse or human gene-specific PCR primers in a 96-well plate using QuantStudio 3 Real-Time PCR system (ThermoFisher Scientific, Rockford, IL). The relative mRNA expressions of the gene of interest were normalized with respect to the housekeeping 18sRNA gene. The data were analyzed using the 2^−ΔΔ*C*^_T_ method and represented as relative fold change.

### Statistical Analysis

The statistical analysis was performed using GraphPad Prism10 V10.1.2 (GraphPad Software, La Jolla, CA). The one-or two-way ANOVA was used to compare the statistical differences between the experimental groups. A P<0.05 was considered statistically significant. The data were expressed as mean ± SD from three biological replicates unless indicated otherwise.

## Acknowledgements

This study is supported by the National Institute of Health (NIH)/ National Eye Institute (NEI) grant R01EY032495 and research start-up funds from the University of Missouri School of Medicine to PKS. The funders had no role in study design, data collection and analysis, decision to publish, or preparation of the manuscript. The Zika virus strain PRVABC59 (NR-50240) was obtained through BEI Resources, NIAID, NIH.

## Authors contributions

ND and PK performed the experiments and analyzed the data. LK helped with fundus and OCT imaging studies. VB is a retina specialist who assisted with clinical scoring of fundus and OCT images as a masked observer. ND and PKS wrote the manuscript. PKS conceived the study, designed and supervised the experiments, and acquired reagents, equipment, and funding for the study. All authors read, edited, and approved the final version of the manuscript.

## Data availability

All relevant data are within the manuscript and figure files.

